# Brief report on the development of patient-derived lung cancer organoids with keratinizing squamous cell carcinoma morphology

**DOI:** 10.64898/2026.04.08.717166

**Authors:** Éabha O’Sullivan, Christina Cahill, Rebecca M O’Brien, Ismail Suliman Elgenaidi, Gavin McManus, William Mc Cormack, Sinéad Hurley, Laura Mary Staunton, Siobhan Nicholson, Stephen P Finn, Ronan Ryan, Gerard J Fitzmaurice, Maeve A Lowery, Jacintha O’Sullivan, Kathy Gately

## Abstract

**Introduction:** Novel therapeutic options are urgently required to improve outcomes and survival for patients with lung squamous cell carcinoma (LUSC). In particular, understanding the unique histological features that define LUSC is essential to improving lung cancer mortality. Many pre-clinical models fail to accurately represent intratumour heterogeneity and recapitulate the tumour microenvironment. This is partly responsible for the poor translation of clinical findings to approved therapies. Our objective was to investigate whether patient-derived organoids, replicate the histological morphological, and structural features of keratinizing LUSC, a poor prognostic subtype of lung cancer.

**Methods:** Organoid cultures were established and maintained from two patients presenting with keratinizing lung squamous cell carcinomas. Immunofluorescent staining of individual organoids and confocal microscopy was performed to confirm expression of tumour markers. Whole organoid domes were fixed, and immunofluorescent staining and imaging was performed to investigate the structural features of the organoid cultures. Findings were compared with histopathological features of the original tumour tissue.

**Results:** Patient-derived organoids expressed tumour markers specific to the squamous cell carcinoma subtype of non-small cell lung cancer, which were confirmed to be expressed in the parent tissue. Within organoid cultures, keratin pearl structures spontaneously developed, matching the keratinizing pattern demonstrated by hematoxylin and eosin staining of the original tumour.

**Conclusions:** Patient-derived organoids have the capability to replicate key histological features of their parent tumour. This high degree of fidelity makes these 3D models an important and valuable tool for understanding complex tumour biology and as a platform for preclinical drug testing to advance novel therapies into the clinic.

## Introduction

Lung squamous cell carcinomas (LUSCs) are associated with high mortality and lack dedicated, disease-specific treatment strategies. Driver mutations commonly detected in lung adenocarcinomas (LUADs) are rarely observed in LUSCs thus targeted interventions have so far provided minimal therapeutic impact^1^. LUSCs often express high levels of the enzyme thymidylate synthase (TS) which limits the effectiveness of pemetrexed-based chemotherapy. Although a subset of LUSCs respond well to immune-checkpoint inhibitors (ICIs) their heterogeneous tumour immune microenvironment (TIME) can foster an immune-evasive niche that is attributed to poor response^2^.

Keratinisation, often a hallmark histopathological feature of well-differentiated LUSC, is associated with impaired immunotherapy efficacy, stemness and a poor prognosis^3,4^. Keratins are the primary intermediate filament proteins in epithelial cells, playing a central role in maintaining structural integrity and protecting cells from physical damage, infection, or xenobiotics^5^. Keratinocytes follow a unique program of terminal differentiation and apoptotic cell death, producing the characteristic keratin layer^6^. In cancer, keratin expression is frequently altered, and high cytokeratin 7 (CK7) expression is associated with increased proliferation, epithelial-mesenchymal transition (EMT), tumour progression, and metastasis^7^. Pharmacological keratin modulation may therefore provide an adjunctive strategy to improve therapeutic efficacy.

Traditional preclinical models often fail to accurately replicate the complex architecture present in human tumours. Patient-derived organoids (PDOs) are three-dimensional (3D) cultures generated from patient tumours that maintain intratumor heterogeneity, preserve key cellular phenotypes, and recapitulate the tumour microenvironment^8^. PDOs have been increasingly applied to study tumour biology, drug response, and personalized therapy. In this study, we established PDOs from patients with keratinizing LUSC, to investigate whether tumour histological features are retained in these 3D models.

## Materials and Methods

### Patient recruitment and patient characteristics

Ethical approval for tumour tissue collection from patients with resectable lung cancer at St. James’s Hospital, Dublin Ireland, was granted by the St. James’s Hospital (SJH) / Tallaght University Hospital (TUH) Joint Research Ethics Committee (JREC) (Submission number 803, Project ID 0723). Patient 1 presented with an 80 mm LUSC of the right lower/middle lobe (cT4N0M0) and underwent pneumonectomy. Patient 2 presented with a 49 mm LUSC of the right upper lobe (cT4N0M0) and underwent lobectomy. Following written informed consent, tumour tissue was sectioned by an experienced histopathologist (SN) and confirmed using haematoxylin and eosin (H&E) staining.

### Generation of PDOs

Fresh tumour tissue was processed within 30 min following resection. PDO generation methodology was adapted from Kim *et al* and Shi *et al*^9,10^. Briefly, tissue was washed, finely minced, and digested while rotating for approx. 1 h at 37°C in DMEM/F-12 (Thermo Fisher Scientific, Waltham, Massachusetts) supplemented with 1% Antibiotic-Antimycotic, 1 mg mL^-1^ collagenase (Sigma-Aldrich, Burlington, Massachusetts), and 0.001% DNAse (Sigma-Aldrich). The dissociated tumour solution was passed through a 100 µm strainer and tumour cells were pelleted, counted, and seeded at 80,000 cells per 50 µL dome in DMEM/F-12 + Cultrex Reduced Growth Factor Basement Membrane Extract, Type 2 (R & D Systems, Minneapolis, Minnesota) (1:10). Cultures were overlaid with specialised growth media (**Supplementary Materials**), maintained at 37°C and 5% CO_2_, imaged regularly, and passaged every 2-3 weeks.

### Immunofluorescence characterisation of organoids

Organoids were gently released from Cultrex domes and fixed with 10% paraformaldehyde (PFA) (Merck, Darmstadt, Germany) for 1 h at 22-25°C. Fixed organoids were collected by gravity sedimentation, washed, and transferred to chamber slides for staining. Samples were permeabilised and blocked (5% FBS, 0.5% Triton-X100) before overnight incubation at 4°C with primary antibodies Phalloidin-iFluor 488 Reagent (Abcam, Cambridge, United Kingdom) and p63-α Antibody (Cell Signalling Technology, Danvers, Massachusetts), followed by further overnight incubation at 4°C with secondary antibody Anti-rabbit IgG (H+L), F(ab’)2 Fragment-Alexa Fluor® 647 Conjugate (Cell Signalling Technology). Following thorough PBS washes, organoids were counterstained with Hoechst 33342 nuclear stain, rinsed, and subsequently imaged using Leica SP8 Scanning Confocal Microscope (Leica, Wetzlar, Germany).

To assess keratinising architecture *in situ*, organoids were fixed and stained within their gel domes, whereby 10% PFA (Merck), permeabilisation/blocking solution, and antibodies were each applied directly to the organoid Cultrex domes. Organoids were stained with Phalloidin-iFluor 488 Reagent (Abcam), Involucrin-Alexa Fluor® Conjugate 647 (Santa Cruz, Dallas, Texas), and Hoechst as before, and whole domes were imaged using Motorised Epifluorescent Microscope. Additionally, organoids were stained with Anti-Human Ki-67 FITC Conjugate (Thermo Fisher), Pan-Cytokeratin (C11)-Alexa Fluor® Conjugate 594 (Santa Cruz), Involucrin-Alexa Fluor® Conjugate 647 (Santa Cruz), and Hoechst, as before, and whole domes were imaged using Leica SP8 Scanning Confocal Microscope.

## Results

### Keratinising lung squamous cell carcinoma PDOs express tumour-specific markers

PDOs were successfully generated from tumour tissue obtained from two individuals with confirmed focally keratinising LUSC (Fig. 1A). Two independent PDO lines, PDO-01 and PDO-02 were derived from patient 1 and patient 2, respectively and were stably maintained in culture, displaying consistent growth characteristics across early passages. Representative bright-field images illustrate the typical morphology of the organoids, which ranged from compact spherical structures to more irregular, multilobular forms (Fig. 1B). To confirm their tumour-specific identity, PDOs were stained for Hoechst nuclear stain, phalloidin cytoskeleton marker, and the squamous lineage marker p63. Robust p63 expression (Fig. 1C) is consistent with the LUSC phenotype and supports that the PDOs faithfully preserve key molecular features of the original tumour.

**Figure 1:**
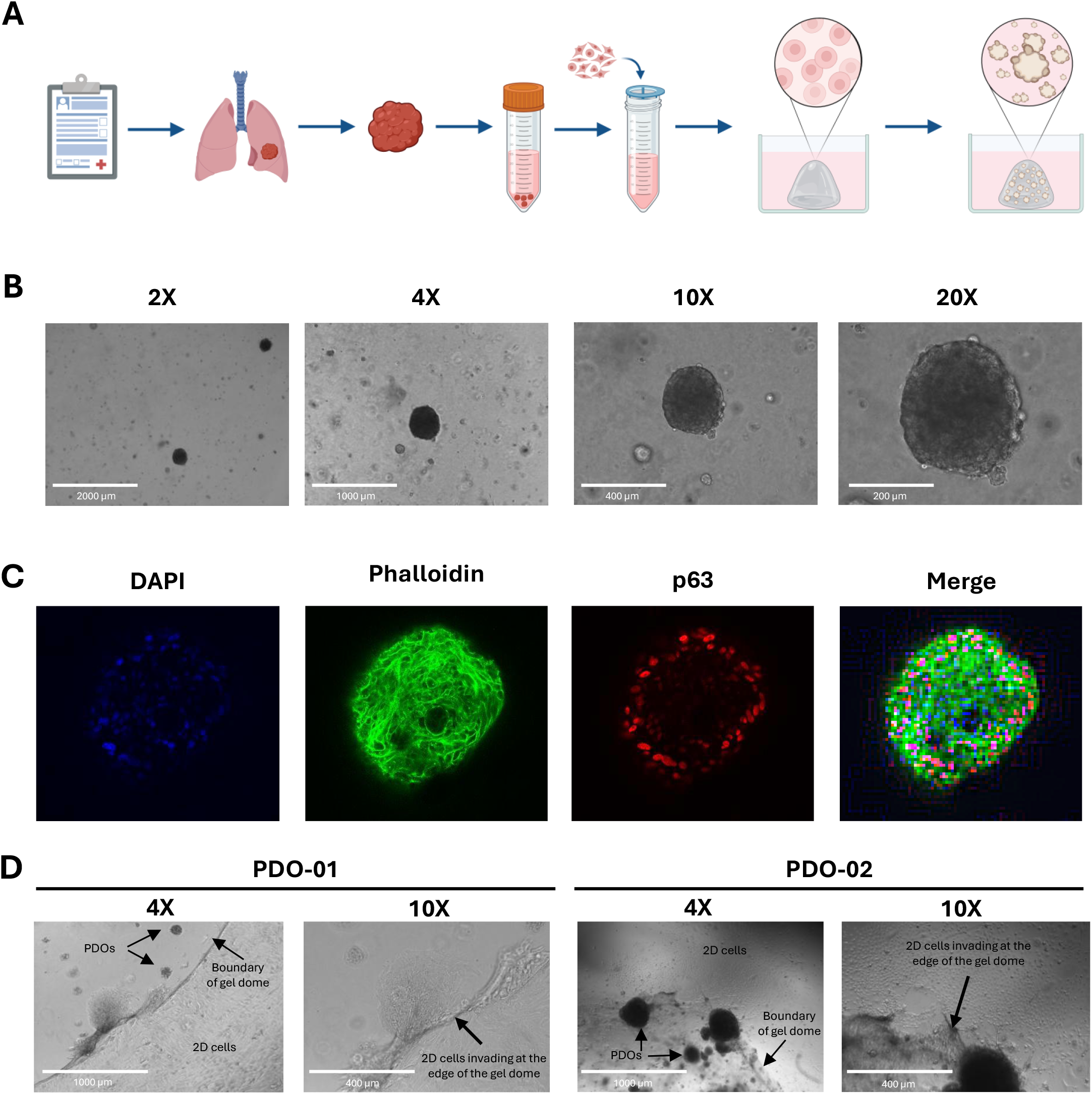
Successful establishment of PDOs from LUSC tissue. **(A)** Resected LUSC tissue was enzymatically dissociated to single cells/clusters, resuspended in Cultrex, and seeded in 50 μL domes. Tumour PDOs were established and maintained in specialised growth media. **(B)** Representative bright field images of established PDOs were captured using an EVOS microscope. **(C)** Immunofluorescent staining was performed of released, fixed PDOs, confirming expression of p63, a LUSC-specific marker. Confocal microscopy was used for imaging. **(D)** Representative bright field images capturing invasive features of LUSC PDOs following 3 passages in culture.

### Lung squamous cell carcinoma PDOs demonstrate invasiveness in vitro

To assess functional behaviour, particularly the invasive potential characteristic of LUSC, PDOs from both patients were observed in culture over time. Following three passages in culture, PDOs began to exhibit invasive features, whereby organoids remained positioned within the gel domes, while distinct populations of adherent 2D cells were observed emerging beyond the boundaries of the matrix, indicative of early invasion (Fig. 1D). Higher-magnification images revealed more detailed features of this invasive front, including elongated cell morphology and progressive migration from the organoid–matrix interface (Fig. 1D). The findings suggest that PDO cultures retain functional tumour behaviours and provide an important platform for studying mechanisms of LUSC invasion and progression.

### Lung squamous cell carcinoma PDOs demonstrate spontaneous keratin pearl formation

To determine whether the PDO cultures reflects the differentiation patterns characteristic of keratinising LUSC, PDOs were observed in culture over time and compared to parent tumour tissue. Examination of the formalin-fixed tumour specimen (Fig. 2A) and its associated H&E-stained section (Fig. 2B) confirmed a focally keratinising squamous phenotype, with well-defined concentric keratin pearls typical of differentiated squamous epithelium. Strikingly, both PDO-01 and PDO-02 recapitulated this hallmark feature *in vitro*. Bright-field imaging (Fig. 2C) revealed circular, laminated structures within the organoids that resembled keratin pearls. These structures were evident across multiple passages and throughout multiple regions of the PDO cultures, appearing spontaneously without induction, indicating that the organoids intrinsically maintain the tumour’s native differentiation hierarchy.

**Figure 2:**
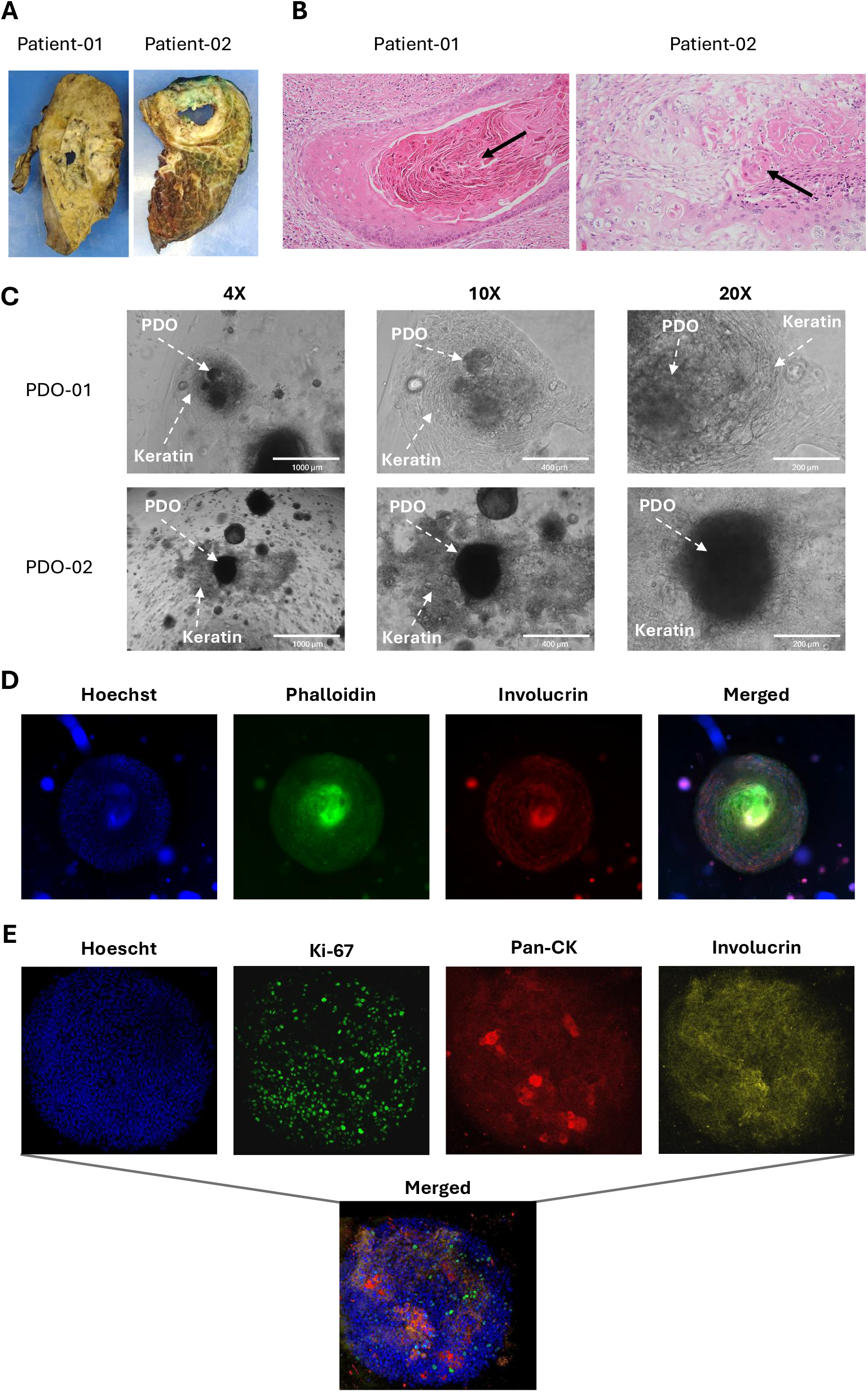
LUSC PDOs recapitulate key histological features of the parent tumour. **(A)** Formalin-fixed LUSC tissue. **(B)** Hematoxylin and eosin staining of parent tissue, demonstrating keratin pearl formation. **(C)** Representative bright field images of LUSC PDOs, captured using an EVOS microscope, demonstrating keratin pearl formation. Immunofluorescent staining was performed of fixed PDOs in gel domes confirming **(D)** expression of Hoechst, phalloidin, and involucrin using epifluorescent microscopy for imaging and **(E)** expression of Hoechst, Ki-67, pan-cytokeratin, and involucrin using confocal microscopy for imaging.

### Keratin proteins are expressed by lung squamous cell carcinoma PDOs

To validate that these structures corresponded to keratinising differentiation, immunofluorescence staining was performed on intact PDOs maintained within gel domes, preserving their three-dimensional organisation and the keratinising structures observed in culture. LUSC PDOs with evidence of keratin pearl–like structures revealed densely populated nuclei throughout the organoid and markedly enriched phalloidin expression in the centre of the organoid, consistent with the formation of a structured, keratinised core (Fig. 2D). Involucrin, a marker of terminal keratinocyte differentiation, was strongly expressed throughout the organoid structure and extended into the surrounding matrix (Fig. 2D).

To confirm these structures reflected true keratin pearl formation, an extended panel of immunofluorescent staining was performed on intact PDOs, demonstrating densely populated nuclei and Ki-67 expression detected broadly throughout the organoid, indicating ongoing proliferative activity consistent with malignant epithelial growth (Fig. 2E). Involucrin expression remained widespread, marking terminally differentiating keratinocytes across the structure. Importantly, cytokeratin staining was positive throughout the organoid and showed intensified signal in discrete regions (Fig. 2E). Together, these findings confirmed that LUSC PDOs not only contained differentiated keratinocytes but also deposited keratin in a manner consistent with the keratinising phenotype of the parent LUSC tumour.

## Discussion

Clinically relevant research models have been the subject of debate for decades. Traditional murine models, while indispensable, often fail to reflect the complexity and heterogeneity of human tumours. Moreover, the time, cost, and labour associated with establishing mouse models limit their translational utility. PDOs offer a compelling alternative, providing a rapid, scalable, and patient-specific platform which can capture the genetic and phenotypic diversity of tumours. These models are particularly valuable for studying tumour subtypes such as keratinising LUSC. In this study, two independent LUSC PDO lines were successfully established from keratinising subtype tumours with the ability to recapitulate tumour markers and spontaneously form characteristic invasive features and keratin pearl structures in PDO cultures.

The ability of PDO models to retain tumour-specific markers ensures that they function as physiologically relevant systems capable of preserving the molecular lineage identity of the primary tumour. The robust expression of p63 strongly verifies the preservation of parent tumour identity as p63 is a defining transcriptional regulator of squamous differentiation and a reliable diagnostic marker for LUSC^11^. Furthermore, broad distribution of Ki-67 positive nuclei, throughout the PDOs, indicates that they also retain the proliferative activity characteristic of malignant squamous epithelium. Ki-67, a key marker of active cell-cycle progression is strongly associated with tumour growth, aggressiveness, and poor prognosis in lung cancer^12,13^. Its widespread expression in this study therefore reinforces that PDOs capture the functional behaviour and the underlying cell-cycle dysregulation of the parent tumour.

Keratinisation and the formation of keratin pearls are features associated with differentiated LUSCs^14^. In an elegant study, Kawai *et al*. investigated the spatial heterogeneity of these structures, which contain peripheral basal-like cells and more central keratinocytes^15^. They demonstrated novel insights into the biology of keratin pearls in LUSC, uncovering a role for NOTCH signalling in the formation of these structures. Recent work has reported keratinising features in LUSC-derived organoids based on histopathological assessment, demonstrating that organoids can recapitulate keratinising tumour morphology observed *in vivo*^16^. Interestingly, we support and extend these findings by demonstrating that keratin pearl–like structures can be grown in vitro directly from dissociated tumour cells and we demonstrate that these structures reflect true keratinising differentiation through immunofluorescent characterisation. Importantly, involucrin, which marks terminally differentiating keratinocytes^17^, and cytokeratin were broadly expressed throughout the PDO structures, in contrast to the central keratinocytes reported by Kawai *et al*. However, centralised phalloidin enrichment was observed in our study, consistent with the presence of actin-rich cytoskeletal organization in differentiating keratinocytes. Additionally, discrete areas of high cytokeratin expression were observed, corresponding to the formation of keratin pearl-like structures, reflecting localized terminal keratinising differentiation, where cells organize into concentric, densely keratinised layers reminiscent of the characteristic keratin pearls seen *in vivo*. These findings parallel well-established histological markers of keratinising squamous differentiation in LUSC and the ability of PDOs to spontaneously reproduce these features indicates that they faithfully capture not only the molecular identity but also the structural and functional heterogeneity of the original tumour.

We highlight that not all LUSC tissue derived from keratinising tumours gives rise to keratinising PDOs, likely reflecting intratumoural heterogeneity and regional variation in keratin-rich areas sampled for PDO generation. Understanding the lung tumour microenvironment *in vivo*, which promotes the induction of keratin pearl formation, and the relationship between tumour progression and keratinisation will support the therapeutic utility of this model. Finally, establishment and maintenance of PDO cultures that recapitulate genetically distinct tumours has the potential to unlock huge molecular and clinical insights and accelerate treatment options for rarer subtypes.

## Data Availability Statement

The data supporting this study are available upon request from the corresponding author.

## Ethical statement

All procedures were performed in compliance with the General Data Protection Regulation (GDPR) guidelines and have been approved by St. James’s Hospital (SJH)/Tallaght University Hospital (TUH) Joint Research Ethics Committee (JREC) and written informed consent was obtained from patients in this study.

## Abbreviations

LUSC: Lung squamous cell carcinoma
LUAD: Lung adenocarcinoma
TS: Thymidylate synthase
ICI: Immune-checkpoint inhibitor
TIME: Tumour immune microenvironment
CK7: Cytokeratin 7
EMT: Epithelial-mesenchymal transition
PDO: Patient-derived organoid
H&E: Haematoxylin and eosin
PFA: Paraformaldehyde

## Author contributions (CRediT statement)

**Éabha O’Sullivan:** Conceptualization, Methodology, Validation, Investigation, Resources, Data Curation, Writing – Original Draft, Visualisation. **Christina Cahill:** Methodology, Validation, Investigation, Resources, Data Curation, Writing – Original Draft, Writing – Review & Editing, Visualisation. **Rebecca M O’Brien:** Conceptualization, Methodology, Validation, Investigation, Resources, Data Curation, Writing – Original Draft, Visualisation. **Ismail Suliman Elgenaidi:** Methodology, Validation, Investigation, Resources, Data Curation. **Gavin McManus:** Software, Formal analysis, Investigation, Resources, Data Curation. **William Mc Cormack:** Conceptualisation, Writing – Review & Editing, Project administration, Funding acquisition. **Sinéad Hurley:** Investigation, Resources, Data Curation, Project administration. **Laura Mary Staunton:** Resources, Data Curation, Project administration. **Siobhan Nicholson:** Investigation, Resources, Writing – Review & Editing. **Stephen P Finn:** Resources, Supervision, Writing – Review & Editing. **Ronan Ryan:** Resources. **Gerard J Fitzmaurice:** Resources. **Maeve A Lowery:** Resources, Project administration, Funding acquisition. **Jacintha O’Sullivan:** Resources, Project administration, Funding acquisition, Writing – Review & Editing. **Kathy Gately:** Conceptualisation, Resources, Writing – Review & Editing, Supervision, Project administration, Funding acquisition.

## Ethical statement

All procedures were performed in compliance with the General Data Protection Regulation (GDPR) guidelines and have been approved by St. James’s Hospital (SJH) / Tallaght University Hospital (TUH) Joint Research Ethics Committee (JREC) (Submission number 803, Project ID 0723, 11^th^ April 2022).

## Acknowledgements

The authors would like to thank the patients and their families for their participation in this study.

## List of supplementary materials

Supplementary Table 1.docx

## References

1. Lau SCM, Pan Y, Velcheti V, Wong KK. Squamous cell lung cancer: Current landscape and future therapeutic options. Cancer Cell 2022; 40(11): 1279–93.

2. Wang Y, Safi M, Hirsch FR, et al. Immunotherapy for advanced-stage squamous cell lung cancer: the state of the art and outstanding questions. Nat Rev Clin Oncol 2025; 22(3): 200–14.

3. Park HJ, Cha YJ, Kim SH, Kim A, Kim EY, Chang YS. Keratinization of Lung Squamous Cell Carcinoma Is Associated with Poor Clinical Outcome. Tuberc Respir Dis (Seoul) 2017; 80(2): 179–86.

4. Zhao J, Shen M, Xu X, et al. Proteomics and single cell profiling identify keratin driven preexisting immunity influences lung squamous carcinoma neoadjuvant therapy. Cancer Lett 2025; 633: 218000.

5. Karantza V. Keratins in health and cancer: more than mere epithelial cell markers. Oncogene 2011; 30(2): 127–38.

6. Candi E, Schmidt R, Melino G. The cornified envelope: a model of cell death in the skin. Nat Rev Mol Cell Biol 2005; 6(4): 328–40.

7. Hosseinalizadeh H, Hussain QM, Poshtchaman Z, et al. Emerging insights into keratin 7 roles in tumor progression and metastasis of cancers. Front Oncol 2023; 13: 1243871.

8. Liu C, Shi C, Wang S, et al. Bridging the gap: how patient-derived lung cancer organoids are transforming personalized medicine. Front Cell Dev Biol 2025; 13: 1554268.

9. Shi R, Radulovich N, Ng C, et al. Organoid Cultures as Preclinical Models of Non-Small Cell Lung Cancer. Clin Cancer Res 2020; 26(5): 1162–74.

10. Kim M, Mun H, Sung CO, et al. Patient-derived lung cancer organoids as in vitro cancer models for therapeutic screening. Nat Commun 2019; 10(1): 3991.

11. Conde E, Angulo B, Redondo P, et al. The use of P63 immunohistochemistry for the identification of squamous cell carcinoma of the lung. PLoS One 2010; 5(8): e12209.

12. Sobecki M, Mrouj K, Colinge J, et al. Cell-Cycle Regulation Accounts for Variability in Ki-67 Expression Levels. Cancer Res 2017; 77(10): 2722–34.

13. Wei DM, Chen WJ, Meng RM, et al. Augmented expression of Ki-67 is correlated with clinicopathological characteristics and prognosis for lung cancer patients: an updated systematic review and meta-analysis with 108 studies and 14,732 patients. Respir Res 2018; 19(1): 150.

14. Drilon A, Rekhtman N, Ladanyi M, Paik P. Squamous-cell carcinomas of the lung: emerging biology, controversies, and the promise of targeted therapy. Lancet Oncol 2012; 13(10): e418–26.

15. Kawai S, Nakano K, Tamai K, et al. Generation of a lung squamous cell carcinoma three-dimensional culture model with keratinizing structures. Sci Rep 2021; 11(1): 24305.

16. Ekanger CT, Ramnefjell MP, Guttormsen MSF, et al. An Organoid Model for Translational Cancer Research Recapitulates Histoarchitecture and Molecular Hallmarks of Non-Small-Cell Lung Cancer. Cancers (Basel) 2025; 17(11).

17. Eckert RL, Welter JF. Transcription factor regulation of epidermal keratinocyte gene expression. Mol Biol Rep 1996; 23(1): 59–70.

