## Supplemental Table 1 for "Brief report on the development of patient-derived lung cancer organoids with keratinizing squamous cell carcinoma morphology"

**Supplementary Table 1: NSCLC PDO media formulation, adapted from Shi et al.^9^**

| **Component** | **Stock**  **Concentration** | **Final**  **Concentration** | **Volume**  **(50mL)** |
| --- | --- | --- | --- |
| Advanced DMEM/F-12 (Thermo Fisher) | - | - | 47,142.5μL |
| B-27 (Thermo Fisher) | 50X | 1X | 1000μL |
| Antibiotic-Antimycotic (Merck) | 10,000u mL^-1^ | 100u mL^-1^ | 500μL |
| GlutaMax (Thermo Fisher) | 200mM | 2mM | 500μL |
| HEPES (Thermo Fisher) | 1M | 10mM | 500μL |
| N-Acetylcysteine amide (NAC) (Merck) | 500mM | 1.25mM | 125μL |
| EGF, recombinant human protein (Bio-Techne) | 100μg mL^-1^ | 50ng mL^-1^ | 25μL |
| FGF-10 (Thermo Fisher) | 100μg mL^-1^ | 100ng mL^-1^ | 50μL |
| FGF-4 (Thermo Fisher) | 100μg mL^-1^ | 100ng mL^-1^ | 50μL |
| A83-01 (Fisher Scientific) | 500μM | 0.5μM | 50μL |
| Y-27632 dihydrochloride (Fisher Scientific) | 10mM | 10μM | 50μL |
| CHIR9902 (Fisher Scientific) | 5mM | 250nM | 2.5μL |
| SAG (Fisher Scientific) | 1mM | 100nM | 5μL |
| **Total Volume** | **-** | **-** | **50mL** |
